# Decoding a cell’s fate: How Notch and Receptor Tyrosine Kinase Signals Specify the Drosophila R7 Photoreceptor

**DOI:** 10.1101/2024.06.23.600273

**Authors:** Ronald A. Arias, Andrew Tomlinson

**Affiliations:** Zuckerman Institute, Department of Genetics and Development, College of Physicians and Surgeons of Columbia University, Jerome L. Greene Science Center, Level 9 Room 028 3227 Broadway New York, NY 10027

**Keywords:** Drosophila R7 photoreceptor, cell fate specification, signal integration, Notch, RTK

## Abstract

The Drosophila R7 photoreceptor is a classic model for understanding how integration of signaling information can provide precise fate directives. It entails extensive interactions between the RTK and Notch signaling pathways, with Notch providing three distinct functions: it both opposes and promotes the general photoreceptor fate, and it determines the specific photoreceptor type. The RTK pathway promotes transcription of *phyl* - a gene expression critical for photoreceptor specification. We show that Notch activity induces transcription of *yan* which encodes a transcriptional repressor of *phyl*. This defines the antagonism between the two pathways, with RTK promoting and Notch opposing *phyl* transcription. We previously showed that Notch provides Sevenless to the cell to allow RTK pathway hyperactivation to overcome the Notch repression, and we now identify the regulation of Yan activity as the integration site of the RTK and Notch functions. Once the cell is specified as a photoreceptor, the third Notch function then prevents *seven-up* (*svp*) transcription. The Svp transcription factor directs the R1/6 photoreceptor fate, and the prevention of its expression ensures the default R7 specification.

**Summary Statement:** This paper examines how the different signals received by the Drosophila R7 photoreceptor precursor are decoded and used to direct the cell’s appropriate differentiation pathway.

## Introduction

Cells in developing tissues can receive signals from nearby or distant cells that direct their fates. The signals can be diffusing peptides or transmembrane proteins presented on neighboring cells and each has a defined receptor type and a corresponding intracellular transduction pathway that it activates. When a cell receives multiple signals, the corresponding transduction pathways are coincidentally active, raising the question of how the information they convey is decoded to direct the correct cell fate. Here, using the developing *Drosophila* eye, we examine how transduction pathways cross-communicate and how these interactions are then manifested as a fate directive for a cell. We focus on the specification of the R7 photoreceptor, which, because of the extensive work previously dedicated to understanding its specification, provides a facile model system of cell fate assignment.

The receptor tyrosine kinase (RTK) and Notch signaling pathways frequently act antagonistically in animal development, with one promoting a specific fate that the other opposes (Sundaram, 2005). But in R7 specification, a more complex interrelationship exists, with Notch activity both antagonizing and promoting the actions of the RTK pathway. This defines the R7 precursor as a cell richly endowed with signaling pathway crosstalk, and it offers itself as an ideal model with which to probe the mechanisms by which multiple signaling pathways are integrated in cell fate specifications.

The fly eye is compound, made from hundreds of subunit ommatidia – each an assembly of 20 different cells (Ready et al., 1976). The R7 photoreceptor is derived from an “equivalence group” of precursor cells that give rise to R7 itself, the R1/R6 photoreceptor pair and the four non-photoreceptor cone cells of each ommatidium. The cells of the equivalence group are assumed to be initially in the same molecular state: expressing the Drosophila EGF Receptor (Schejter and Shilo, 1989) (DER –an RTK) and Notch (N – the founding member of the Notch superfamily of receptors) on their plasma membranes and the Tramtrack (Ttk88) transcription factor in their nuclei (Xiong and Montell, 1993) (Fig1.F). The differential receipt of the RTK and N signals then determines the individual fates of these cells.

The ligand for DER is Spitz (Schweitzer et al., 1995), a peptide that diffuses and activates the RTK pathway in all R7, R1/R6, and cone cell precursors (Freeman, 1996) (Fig.1A-C green arrows). The N ligand, Delta (Dl), is expressed by the R1/6 precursors as they begin their photoreceptor differentiation (Tomlinson and Struhl, 2001), and this activates N in the R7 and cone cell precursors. Thus, in R1/6 precursors, only DER is active, whereas in the R7 and cone cell precursors, both pathways operate. Ttk represses the neural fate, and in those cells destined to be photoreceptors (R1/6/7), it is removed by the action of the RTK pathway (Li et al., 1997, Li et al., 2002). But in the cone cell precursors, Ttk persists, and these cells are specified as non-neuronal support cells. In directing the R7 fate, N plays three distinct functions (Tomlinson et al., 2011). First, in its *anti-neural function*, it acts to prevent the DER pathway from removing Ttk in R7 and cone cell precursors. Second, in its *neural-promoting function*, it supplies Sevenless (Sev - another RTK (Hafen et al., 1987)) to these cells, which is selectively activated in the R7 precursor and not in the cone cells. Sev provides an RTK pathway hyperactivation that overcomes the N *anti-neural function* resulting in Ttk degradation. Third, in its *R7-v-R1/6 function,* N then acts in R7 precursors to ensure that it differentiates as the specialized R7 photoreceptor rather than the generic R1/6 type.

**Figure 1.**
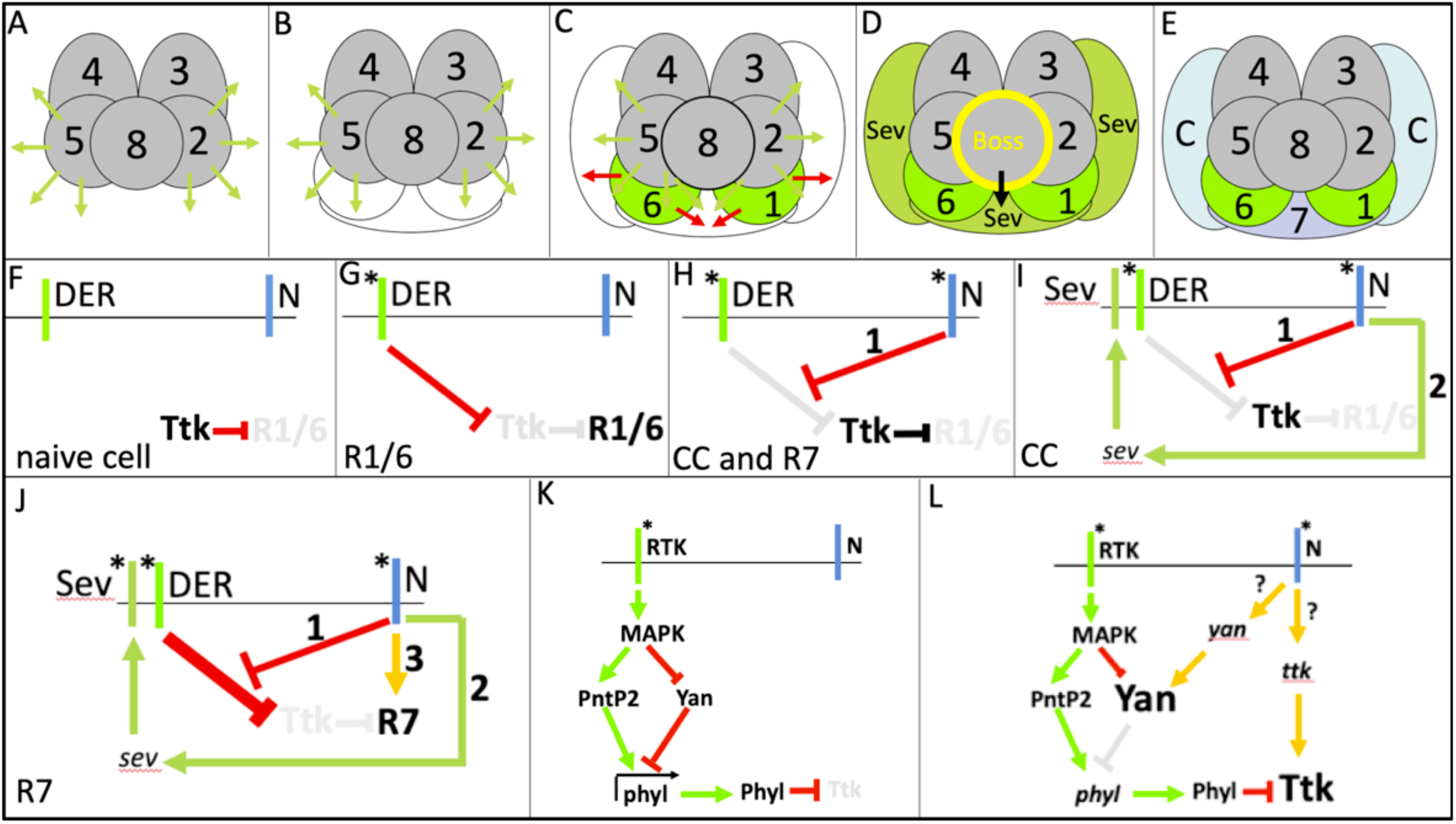
Schematic diagrams of the cells of the developing ommatidia, the signals they send and receive, and the interactions between them. **(A-E)** The cell patterning and signaling of developing ommatidia. **(A)** The precluster (R2,3,4,5,and 8) is a discrete grouping of cells from which the R2/5 pair are thought to release the DER ligand Spitz (green arrows). **(B)** The precursors of the R1/6/7 (white profiles) are added to the R2/8/5 face of the cluster. Spitz signal reaches the cells in the R1/6 position first. **(C)** The receipt of Spitz triggers the R1/6 precursors to begin differentiation (green shading) as two cone cell precursors are added to the flanks of the structure. The differentiating R1/6 cells express Dl (red arrows) that activates N in the R7 and cone cell precursors. **(D)** N activity in R7 and cone cell precursors leads to their expression of Sev (darker green). Boss, the Sev ligand, is membrane-tethered and is expressed exclusively by R8. The R7 precursor contacts R8 and its Sev is activated. However, the cone cell precursors are distant from R8 and their Sev remains inactive. **(E)** Sev activation leads to the specification of the R7 precursor whereas Sev inactivity in the other cells leads to their specification as cone cells. (F-J) Summaries of the RTK and N interactions in the various cells. **(F)** Prior to incorporation into the cluster, the naïve cells express the DER and N receptors and the transcription factor Ttk, which represses the photoreceptor fate. **(G)** In R1/6 precursors, DER activation (asterisk) leads to the removal of Ttk, allowing the specification of the photoreceptor fate. The absence of N activity leads to the default R1/6 type. **(H)** In the R7 and cone cell precursors, N is additionally active, and one function of N here (1) is to prevent DER activity from removing Ttk. **(I)** N activity also promotes Sev expression (2) in R7 and cone cell precursors. **(J)** In R7 precursors (but not cone cells), Sev is activated (asterisk). The hyperactivation of the RTK pathway overcomes the repression to Ttk removal and the cell is specified as a photoreceptor. N activity (3) supplies the information that directs the R7 rather than R1/6 fate. **(K,L)** Details of how the RTK pathway promotes *phyl* transcription and the two potential targets for N (question marks) in opposing it. **(K)** Acting through MAPK, the RTK pathway inactivates Yan, the repressor of phyl transcription, and potentiates the ability of the PntP2 to promote it. **(L)** We considered two gene targets of N that could oppose the ability of the RTK pathway to remove Ttk. If N activity promotes yan transcription and leads to increased Yan proteins levels, this would titrate the actions of the RTK protein to inactivate it. Ifr on the other hand, N activity promotes ttk transcription, then the increased levels of Ttk proteins would titrate the effects of Phyl in degrading it.

In R7 precursors, both the *N anti-neural* and *N neural-promoting* functions are active, with the former opposing the photoreceptor fate (countering Ttk removal) while the other, through the action of Sev, overrides that opposition. This poses the question of how these opposing functions are integrated in the R7 precursor; what are their molecular targets, and how do they collectively lead to the removal of Ttk? A key transcriptional target of the RTK pathway in equivalence group cells is the gene *phylopod* (*phyl),* the expression of which directs the removal of Ttk. Prior to the activation of the RTK pathway, *phyl* transcription is opposed by the direct action of the Yan ETS transcriptional repressor protein. RTK signaling activates MAPK, which phosphorylates and inactivates Yan, thereby relieving the block to *phyl* transcription. Additionally, MAPK phosphorylates and activates PointedP2 (PntP2 –an ETS transcriptional activator), which then replaces Yan at the *phyl* locus to drive transcription (Brunner et al., 1994, O’Neill et al., 1994, Lai and Rubin, 1992). The resulting Phyl protein then directs the removal of Ttk (Fig.1K). This defines a simple switch role for the RTK pathway – it acts to promote *phyl* transcription and thereby species the photoreceptor fate. In R1/R6 precursors (in which N is inactive), a moderate RTK activation suffices for this purpose, but in the R7 precursor (in which the *N anti-neural function* is present), a Sev-mediated hyperactivation is necessary. In this paper, we show that N activity in the R7 precursor raises *yan* transcription, and we posit that this represents the *N anti-neural function*, with the resulting higher level of Yan protein overcoming the ability of MAPK to inactivate it and leaving a functional repressor complex at the *phyl* locus. Consistent with this model, we observe a corresponding increase in R7 *phyl* transcription resulting from the loss of N activity. We therefore identify transcriptional regulation of *yan* and *phyl* as key points of integration of the RTK and N pathways in the removal, or not, of Ttk. Hence, we propose that the N *anti-neural* function opposes the transcription of *phyl* while the Sev-mediated *neural-promoting function* hyperactivates the RTK pathway to negate this (by inactivating Yan) and actively promotes *phyl* transcription through the agency of PntP2.

The *R7-v-R1/R6 function* of Notch comes into play after the removal of Ttk from R7 precursor and operates to define it as the R7 versus the R1/R6 type of photoreceptor. The Seven-up (Svp) transcription factor has long been associated with this function. It is expressed in R1/R6 but not R7 precursors, and when its expression is lost from R1/R6 precursors, they transform into R7-like cells. Conversely, when Svp is ectopically expressed in R7 precursors they transform into R1/R6-like cells (Mlodzik et al., 1990, Mlodzik et al., 1992, Hiromi et al., 1993, Miller et al., 2008). Thus, in cells of the R7 equivalence group, if a cell removes Ttk and does not express Svp, it becomes an R7, but if it does express Svp, it is specified as an R1/R6 type. Here we show that N activity prevents *svp* transcription in the R7 precursor, thereby preventing any Svp expression and ensuring the R7 photoreceptor fate. Additionally, we show that when R1/6 precursors are re-specified as cone cells (by blocking their removal of Ttk), they persist in their N-dependent Svp expression, arguing that Svp expression does not require the removal of Ttk. We thus view two sequential steps in the specification of the R7 cell. First is the integration of the *N anti-neural* and *neural-promoting functions* that achieve Ttk removal and the loss of the repression of the photoreceptor fate. Second, and independently of that integration, the N *R7-v-R1/6 function* prevents Svp expression in the R7 precursor that ensures that it adopts the appropriate photoreceptor fate.

## Results

### Overview

In this results section, we address the nature of the N *anti-neural* and *R7-v-R1/6 functions* in the R7 precursor. First, we describe a new tool that selectively abrogates N activity in R7 precursors. We then examine the activity transcriptional reporters for candidate N target genes (whether direct or indirect) for the *anti-neural* and *R7-v-R1/6 functions* and determine whether their expression levels in R7 precursors changes when their N activity is compromised.

### A new method for abrogating N activity in the R7 precursor

*N* clones do not survive in the developing eye, making it difficult to assess their frank loss of function. Previously, we used a variety of techniques to down regulate N activity, but each was problematical for a variety of reasons. Here, we describe a simple, facile technique that effectively abrogates N activity selectively in the R7 precursor. The *scaG109-Gal4* line is expressed in R8s in ommatidia of the eye disc (Fig. 2A shows a UAS.GFP reporter of its activity). In early ommatidia, the expression is evident in R8s, but other cells may stain (usually R2/5 – arrows Fig.2B). However, as the ommatidia mature, expression becomes restricted to R8s. The reporter levels in the more mature R8s vary in intensity with occasional examples where they are reduced or absent (Fig.2B white and red circles respectively). When *scaG109-Gal4* was crossed into a *UAS.Dl-Dominant-Negative* (*DlDN* – a form of Dl lacking the majority of its intracellular domain) background (hereafter referred to as *R8-DlDN*) and examined in the adult eyes, the vast majority of cells in the R7 positions differentiated as R1/6-like cells. Fig 2C shows *wild type* ommatidia in which the small central R7 rhabdomere is evident (red arrow), but in *R8-DlDN* (Fig.2D) the cells in the R7 positions show large peripheral rhabdomeres characteristic of the R1/6 photoreceptors (back arrow). To confirm that these R1/6-like cells in the R7 positions indeed derive from R7 precursors, we examined developing eye discs. In *wild type* ommatidia, cells in the R7 position express the Runt and Elav transcription factors but do not show the Svp marker characteristic of the R1/6 type (Fig. 2F). In *R8-DlDN* eye discs, the cell in the R7 positions do not express Runt but show Elav and Svp, consistent with their transformation into R1/6 types (Fig. 2G). R1/6 precursors do not require Sev for their specification, and when we examined *R8-DlDN* in a *sev°* background, we observed that the cells in the R7 positions still appeared as R1/6 types (Fig. 2E arrows). To evaluate the levels of N activity in these *R8-DlDN* R7 precursors, we examined expression of two known N response genes, *sev* and *deadpan (dpn)*. We previously showed that R7 *sev.lacZ* expression (Fig.2I) is lost when N activity is compromised in R7 precursors (Tomlinson et al., 2011), and in the *R8-DlDN* background, R7s also displayed the loss of *sev.lacZ* expression (Fig.2J’). Fig.2K shows the expression of Dpn protein in a wild type eye disc. There are two bands of nuclear staining. The black line indicates the expression of Dpn in early ommatidia, and the red line highlights the position of R7 Dpn expression. Strikingly, in the presence of *R8-DlDN*, the vast majority of cells in the R7 positions do not express Dpn. The white circle indicates two ommatidia in which Dpn expression is not lost, which likely correspond to the rare ommatidia in which *scaG109-Gal4* expression is stochastically reduced or absent. Thus, not only does *R8-DlDN* cause cells in the R7 position to develop as R1/6-like cells, it also knocks down expression of two known N response reporters. We infer that it acts to potently reduce N activity in the R7 precursor. N activation in the R7 precursor is triggered by the expression of Dl by it R1 and R6 neighbors, and we surmise that the over-expression of DlDN by R8 sequesters N on the R7 plasma membrane into inactive complexes and prevents normal N pathway activation.

**Figure 2.**
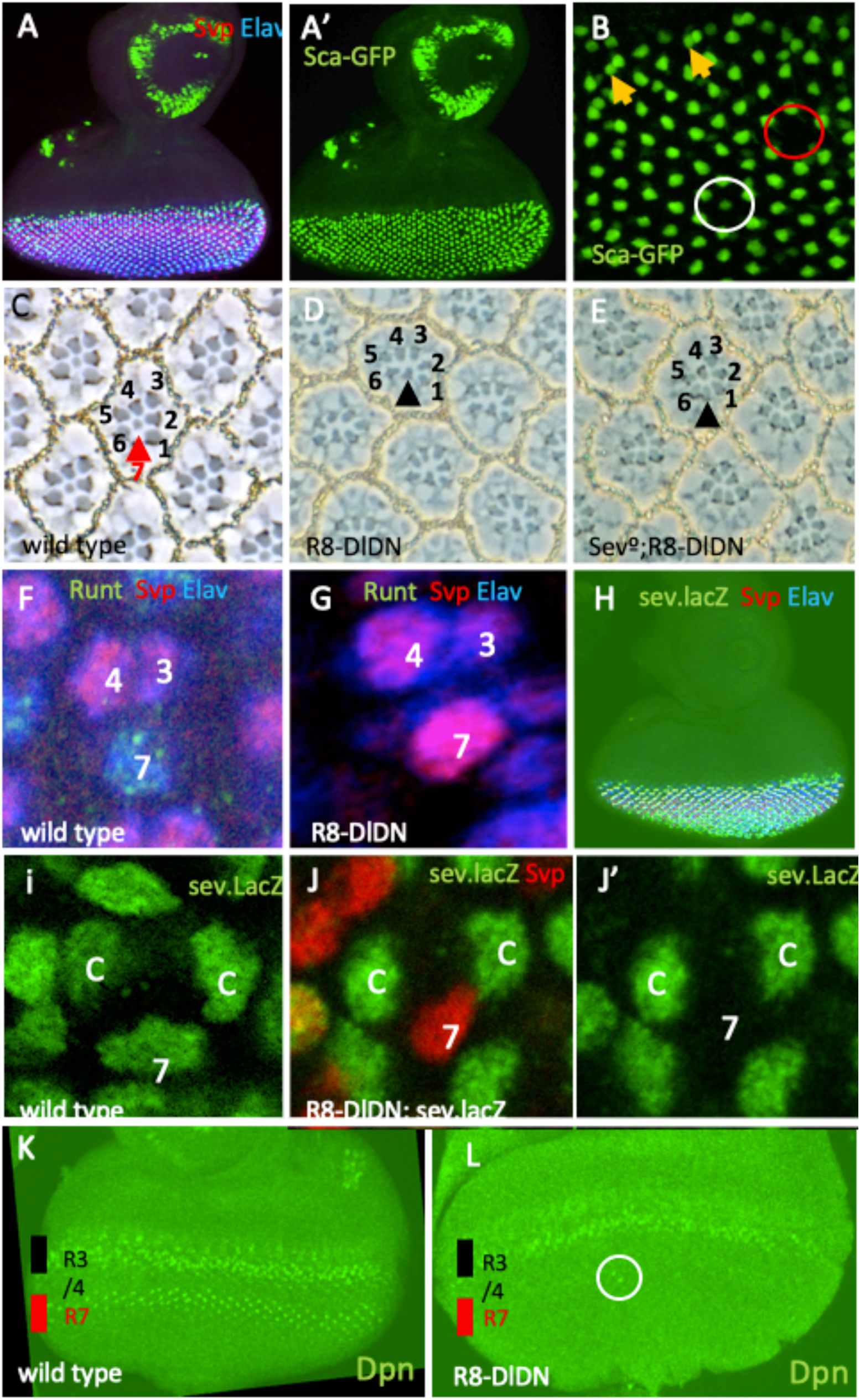
R8-DlDN selectively compromises N activity in the R7 precursor. (**A, A’**) When *sca G109 Gal4* drives *UAS-GFP* expression (*sca-GFP*), GFP expression is detected in many tissues in the eye antennal disc. (**B**) Behind the furrow in the eye disc, expression is observed in R8 cells. Expression in R8-adjacent cells (usually R2/5) is seen particularly in younger ommatidia (yellow arrows). Infrequently, the GFP level in R8 is reduced (white circle) or absent (red circle). (**C-E**) Sections through adult eyes. (**C**) In wild type ommatidia, the cell in the R7 position had a small central rhabdomere (red arrow). (**D**) In an *R8-DlDN* background, the vast majority of cells in the R7 positions appears as R1/6 type photoreceptors (black arrow). (**E**) In a *sev° R8-DlDN* background, the vast majority of cells in the R7 positions appears as R1/6 type photoreceptors (black arrow). (**F-L**) Images from third instar eye discs in various genetic backgrounds. (**F**) In wild type ommatidia, the cell in the R7 position expresses Elav and Runt but not Svp (**G**). In *R8-DlDN* discs, the cell in the R7 position expresses Svp instead of Runt, indicating a transformation into the R1/6 type. (**H**) Image of an eye-antennal disc expressing *sev.lacZ* and counterstained for Svp and Elav. (**I**) In wild type, *sev.lacZ* is expressed in the R7 precursor along with cone cell precursors. (J, J’) In an *R8-DlDN* background the cell in the R7 position expresses Svp but not *sev.lacZ* (compare I with J’). (**K, L**) Eye discs stained for Dpn. (**K**) In wild type, staining is evident in two bands. The upper band (black line) indicates Dpn expression in preclusters, and the lower band (red line) highlights the expression of Dpn in the R7 precursors. (**L**) In *R8-DlDN* discs, the precluster Dpn staining is normal, but in the vast majority of the R7 precursors, it is absent (the white circle indicates two ommatidia in which R7 staining is evident).

### Integration of the N *anti-neural* and *proneural functions* in R7 specification

Our goal is to understand how the information from the N and RTK pathways is integrated in the R7 precursor to specify the R7 fate and to define the genes that are regulated by the N functions. In this section, we examine the genes that N regulates to execute its *anti-neural function* to understand how the block to photoreceptor specification (opposition to Ttk removal) is imposed.

### *yan* appears as an R7 N target gene

MAPK plays a key role in mediating the RTK signal that promotes the photoreceptor fate by inactivating the Yan repressor and activating the PntP2 transcriptional activator to raise *phyl* transcription levels (Fig.1L). The *yan* gene appeared as a potential target of the N *anti-neural function* for two reasons. First, its transcription is known to be regulated by N activity in eye discs (Rohrbaugh et al., 2002), making it a likely target in R7 specifically. Second, if N activity promotes *yan* transcription in R7, this likely leads to increased Yan protein levels which may titrate the effects of MAPK to phosphorylate and inactivate it. We used Crispr to knock in the GFP^NLS^ coding sequence into the *yan* locus to produce a transcriptional reporter (Fig.3). This shows that *yan* is widely transcribed in imaginal tissues (Fig.3A-D). In the eye-antennal disc, it is ubiquitously expressed with a pronounced upregulation in the eye field (Fig.3A,C,D) with evident staining of the nuclei in the basal region belonging to the cells that will be later recruited to the ommatidia (Fig.3D). As nuclei rise from this basal pool to join the ommatidial clusters, some cells show an upregulation of expression. The details of these expressions are given in the legend to Fig.3, but the relevant cells here are R7 and cone cell precursors. As the presumptive R7 nucleus rises, it shows a burst of GFP expression (Fig. 3G) which later declines (not shown). Subsequently, the cone cell nuclei rise and show a persistent increased expression (Fig.3H). We crossed the *yan* reporter into the *R8-DlDN* background and observed the cell in the R7 position with compromised N activity (as evidenced by the expression of Svp) (Fig.3I), with a corresponding loss of the burst of GFP expression (compare Fig.3G and I’). Thus, we detect a burst of *yan* transcriptional activity that is lost when N activity is abrogated, which is consistent with a role in mediating the N *anti-neural* function. When the *yan* reporter was examined in a *sev°* background (Fig.3J,J’), the burst of GFP expression was unchanged, but the later decline did not occur (consistent with the transformation into a cone cell precursor), suggesting that the Sev-dependent hyperactivation of the RTK pathway has no influence on the initial burst of N-dependent *yan* transcription.

**Figure 3.**
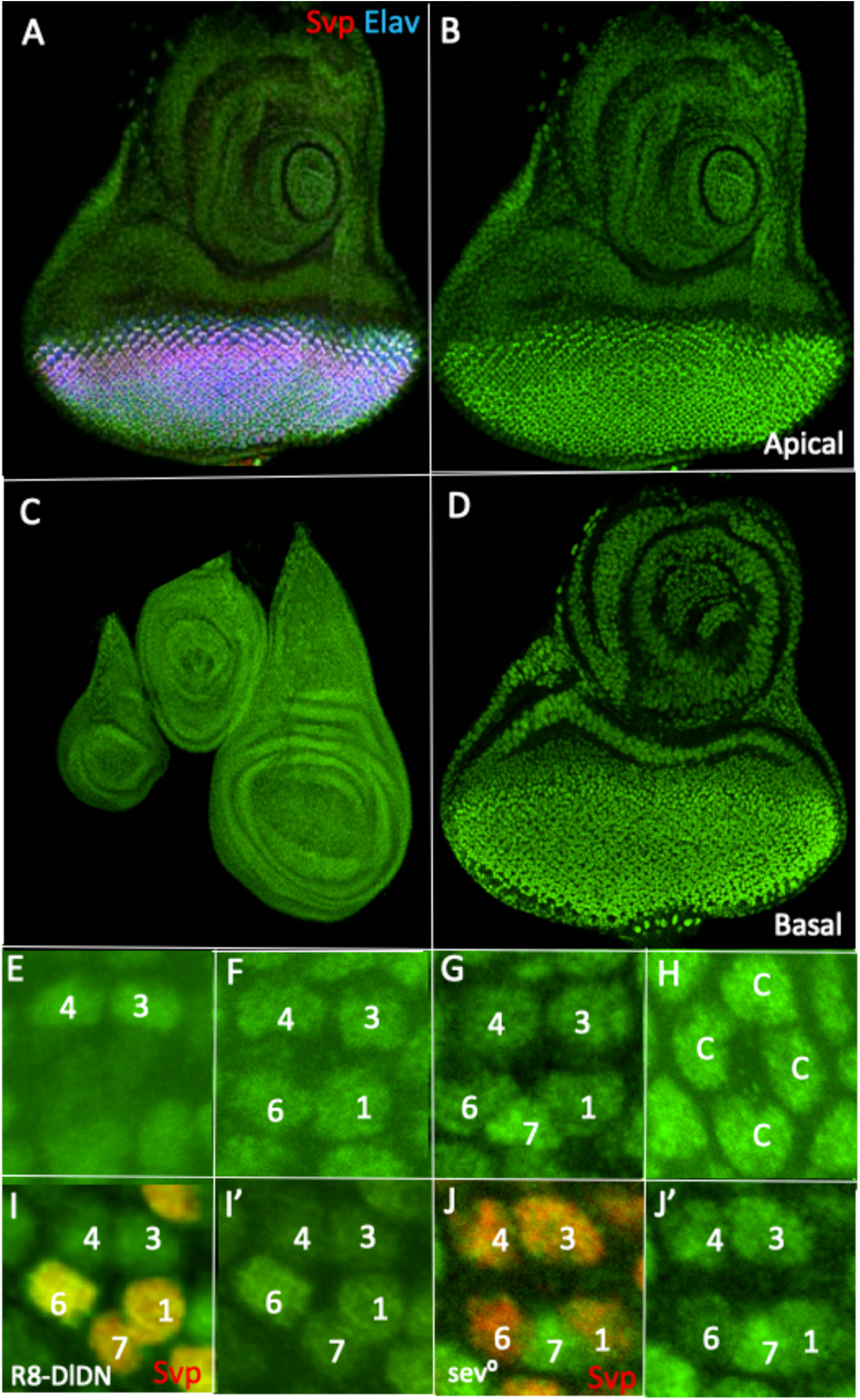
Expression patterns of *yan.GFP* under various genetic conditions. (**A,B**) *yan.GFP* expression in a *wild type* eye antennal disc show an upregulation behind the furrow that precedes the differentiation of any photoreceptor as evidenced by Elav and Svp expression. (**C**) *yan.GFP* in a wing, haltere, and leg disc indicate that the gene is widely expressed in imaginal tissue. (**D**) *yan.GFP* expression shows extensive basal expression in the eye region behind the furrow. (**E-H**) *yan.GFP* expression in maturing ommatidia. (**G**) There is a burst of expression in the R7 precursor which subsequently declines. (**H**) Cone cells show strong and persistent *yan.GFP* expression. (**I,I’**) When N activity is compromised in the R7 precursor as evidenced by its expression of Svp, there is a corresponding loss of the burst of *yan.GFP* in the R7 precursor (compare R7 expression in G and I’). (**J,J’**) In *sev°*, the cell in the R7 position shows high-level persistent *yan.GFP* expression.

### *ttk* does not appears as an R7 N target gene

In Drosophila sensory organ precursors, a Phyl-dependent removal of Ttk occurs in a manner similar to that which happens in the R7 precursor (Pi et al., 2004, Pi et al., 2001), and since N was thought to raise *ttk* transcription, we considered it a potential target for the N *anti-neural function*. That is, if *ttk* transcription is raised by N activity, then the increased Ttk protein could titrate the efforts of Phyl and its associated proteins in degrading it leading to persistent Ttk and the repression of the photoreceptor fate (Fig.1L) We therefore generated a ttk.GFP^NLS^ transcriptional reporter (Fig.4). It showed extensive expression in imaginal tissue (Fig.4A-C), and in the eye antennal disc, the eye region showed low-level expression in the basal pool of cells, which was maintained in the photoreceptors nuclei as they were recruited to the clusters and rose apically. In cone cell precursors a pronounced increase in GFP expression was detected. In the R7 precursors, the level of GFP expression levels was the same as the other photoreceptors and significantly less than that seen in the cone cells (Fig.4D). When examined in the *R8-DlDN* background, the cell in the R7 positions with abrogated N function (as evidenced by its Svp expression (Fig 4F)) showed no detectable difference in GFP expression from its wild type counterpart (compare Fig. 3D and F’). When examined in *sev°*, the R7 precursor showed cone cell levels of expression (Fig.4G). This result was expected because *sev* function is required for Dpn expression, which then reduces *ttk* expression levels in the R7 precursor (Mavromatakis and Tomlinson, 2016).

**Figure 4.**
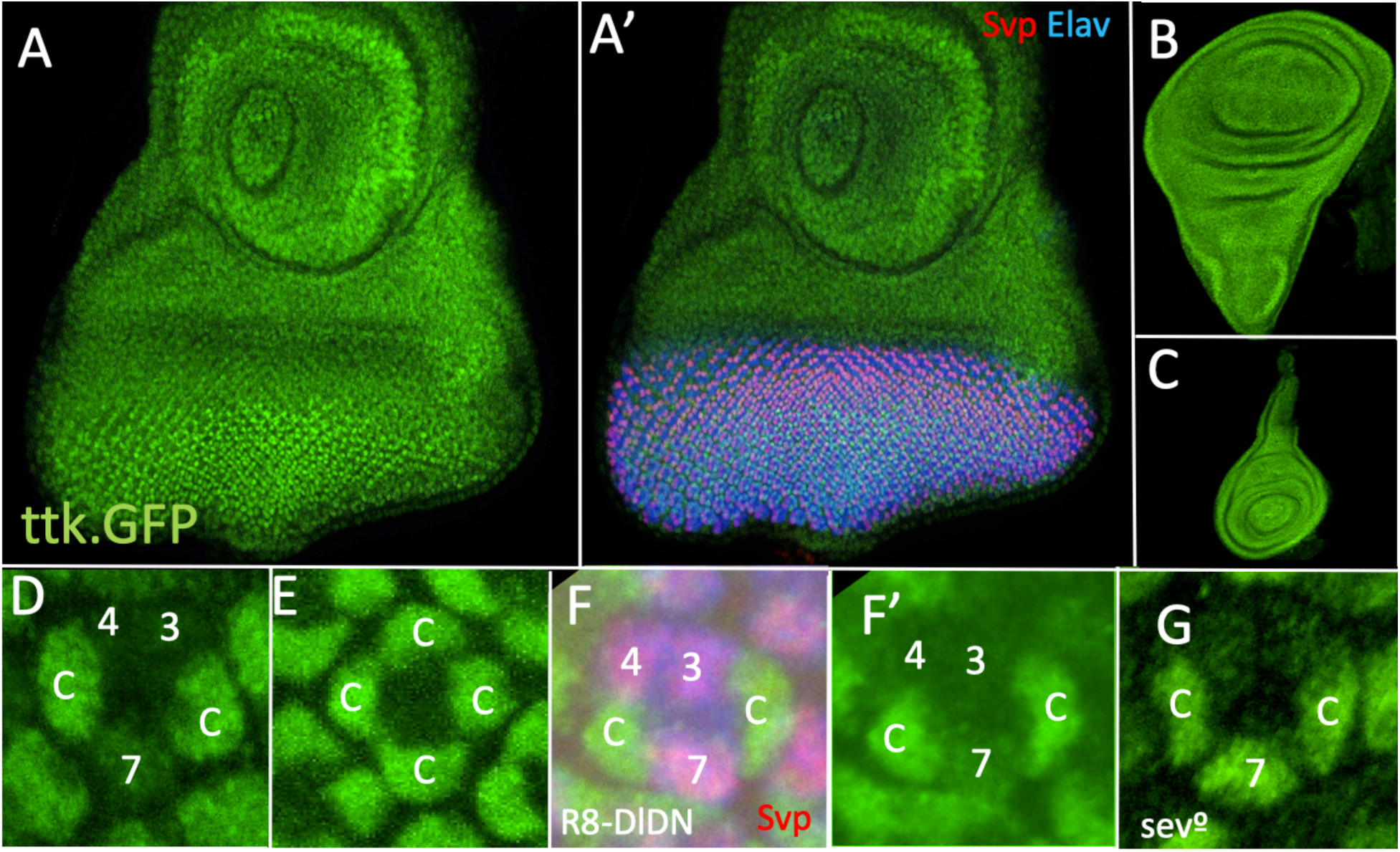
Expression patterns of *ttk.GFP* under various genetic conditions. (**A-C**) *ttk.GFP* is expressed widely in imaginal tissues and is upregulated in the eye disc behind the furrow. (**D,E**) *ttk.GFP* expression is low in R7 precursors but is upregulated in cone cell precursors. (**F, F’**) When N activity is compromised in the R7 precursor (*R8-DlDN*) evidenced by Svp expression, low levels of *ttk.GFP* are present. (**G**) When *sev* gene function is removed (*sev°*), *ttk.GFP* expression levels in the R7 precursor appears as in cone cell precursors.

### *phyl* transcription is negatively regulated by N activity

The results above suggest that N activity triggers a burst of *yan* transcription in the R7 precursor which later declines. If this leads to a meaningful increase in Yan protein that can effectively titrate the actions of MAPK to inactivate it, and since Yan acts to repress *phyl* transcription, then we expect a corresponding N-dependent downregulation of *phyl* expression caused by persistent transcriptional repression by Yan. To evaluate this, we generated a *phyl.GFP^NLS^*transcriptional reporter and observed expression in imaginal tissue in regions fated to generate sensory organs (Fig.5A-C). In the eye, expression is detected in the furrow preceding the formation of the preclusters - Fig.5A’ shows GFP expression ahead (anterior) of the Elav and Svp cluster staining. GFP expression is maintained in the basal pool of cells, and as the nuclei of the prospective photoreceptor nuclei rise, expression levels increase with enhanced levels in R3/4 (Fig. 5B - the expression in R8 subsequently rise as the clusters mature (not shown)). High levels are then seen in R1/6 (Fig.5E) and then in R7 (Fig.5F), but as the cone cells join, they maintain the levels of the basal pool and show considerably weaker levels of expression than the photoreceptors (Fig.5G). We then crossed the *phyl* reporter into the *R8-DlDN* background and observed no change in GFP levels in the R7 precursor (compare Fig.5 F and H’). Naively, we might interpret this as evidence that N activity does not regulate *phyl* transcription, but it is important to note that abrogation of N activity also removes *sev* transcription (Fig.2J). We therefore crossed the *phyl* reporter into a *sev°* background and observed a reduction of expression in the R7 precursor to the levels normal for cone cells (compare Fig.5I’ and F). In this *sev°* background, we infer the continued presence of the N repression, and when we compare the loss of Sev (*sev°*) with the joint loss of Sev and the repression (*R8-DlDN*), we detect the N-mediated downregulation of *phyl* transcription (compare Fig.3H’ and I’). We next examined whether restoring Sev (in a N-independent manner using *GMR.sev*) in an *R8-DlDN* background would hyperactivate *phyl* transcription in R7s because the Sev-dependent positive input would no longer be opposed by the repression. This we did not observe; in *R8-DlDN; GMR.sev* eye discs, transcription of *phyl* appeared at wild type levels in R7 precursors (Fig.5K’). There are many reasons why we might not observe hyperactivation of *phyl* transcription in *DlDN;GMR.sev* R7 precursors, which we address in the Discussion along with a review of the reasoning that *phyl* transcription is downregulated by N activity. Lastly, we examined the *phyl* reporter in the *sev°;R8-DlDN* double mutant and, as expected, observed expression levels typical of *R8-DlDN* alone (compare Fig.5H’ with K’).

**Figure 5.**
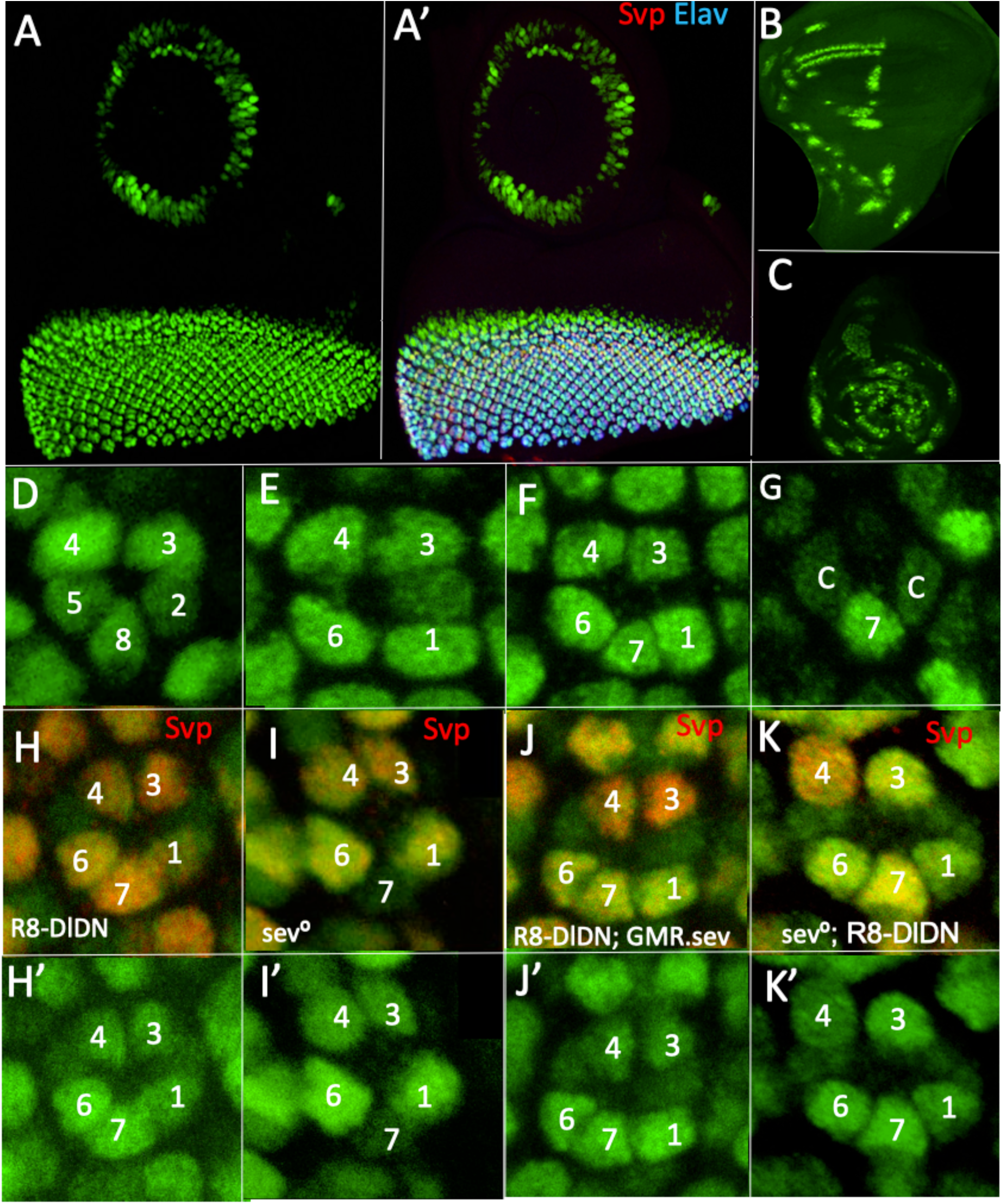
Expression patterns of *phyl.GFP* under various genetic conditions. (**A, A’**) *phyl.GFP* expression in a *wild type* eye antennal disc show an upregulation behind the furrow that precedes the differentiation of any photoreceptor as evidenced by Elav and Svp expression. (**B**) Wing discs and (**C**) leg discs show *phyl.GFP* expression in sensory organ precursors. (**D-G**) *phyl.GFP* expression in maturing ommatidia; (**F**) expression is strong in the R7 precursor and (**G**) remains so as the cone cell precursors show significantly reduced expression levels. (**H, H’**) In *R8-DlDN* clusters, R7 expresses Svp and show normal levels of *phyl.GFP* expression (compare F with H’). In *sev°*, there is a reduction in R7 *phyl.GFP* expression to that seen in the cone cells (compare G with I’). (**J, J’**) When N activity is compromised in the R7 precursor (*R8-DlDN*) and Sev expression is restored to the cell (*GMR.sev), phyl.GFP* expression levels remain normal. (**K, K’**) When N activity is compromised in the R7 precursor (*R8-DlDN*) and *sev* gene function is removed (*sev°*), normal levels of *phyl.GFP* are observed.

### Regulation of N R7-v-R1/6 function

The competition between N *anti-neural* and *neural-promoting functions* determine whether an equivalence group cell is specified as a photoreceptor or a cone cell. In those destined to become photoreceptors, an additional N activity (the *R7-v-R1/6 function*) then determines whether a cell is specified as an R7 or R1/6 type (Fig.1J yellow arrow labeled 3). Svp is a steroid receptor-like transcription factor that is expressed in the R1/6 cells of the R7 equivalence group but not in R7 or cone cell precursors (Mlodzik et al., 1990). In its absence, R1/6 cells transform into R7-like photoreceptors (Mlodzik et al., 1990, Miller et al., 2008), and when Svp is ectopically expressed in R7 precursors, they are respecified as R1/6-like cells (Hiromi et al., 1993). Thus, Svp appears as a candidate transcription factor that mediates the N *R7-v-R1/6 function*. Svp expression has long been suspected to be repressed by N (Miller et al., 2008), and here we used the *svp.lacZ* line to evaluate this at the transcriptional level. *svp.lacZ* is expressed only in the R3/4 and R1/6 precursors and is specifically absent from R7 (highlighted by Dapi – Fig. 6A). In the R8-DLDN background, *sev.lacZ* was ectopically expressed in the R7 precursor (Fig.6B), indicating that N functions to repress *svp* transcription in the cell.

**Figure 6.**
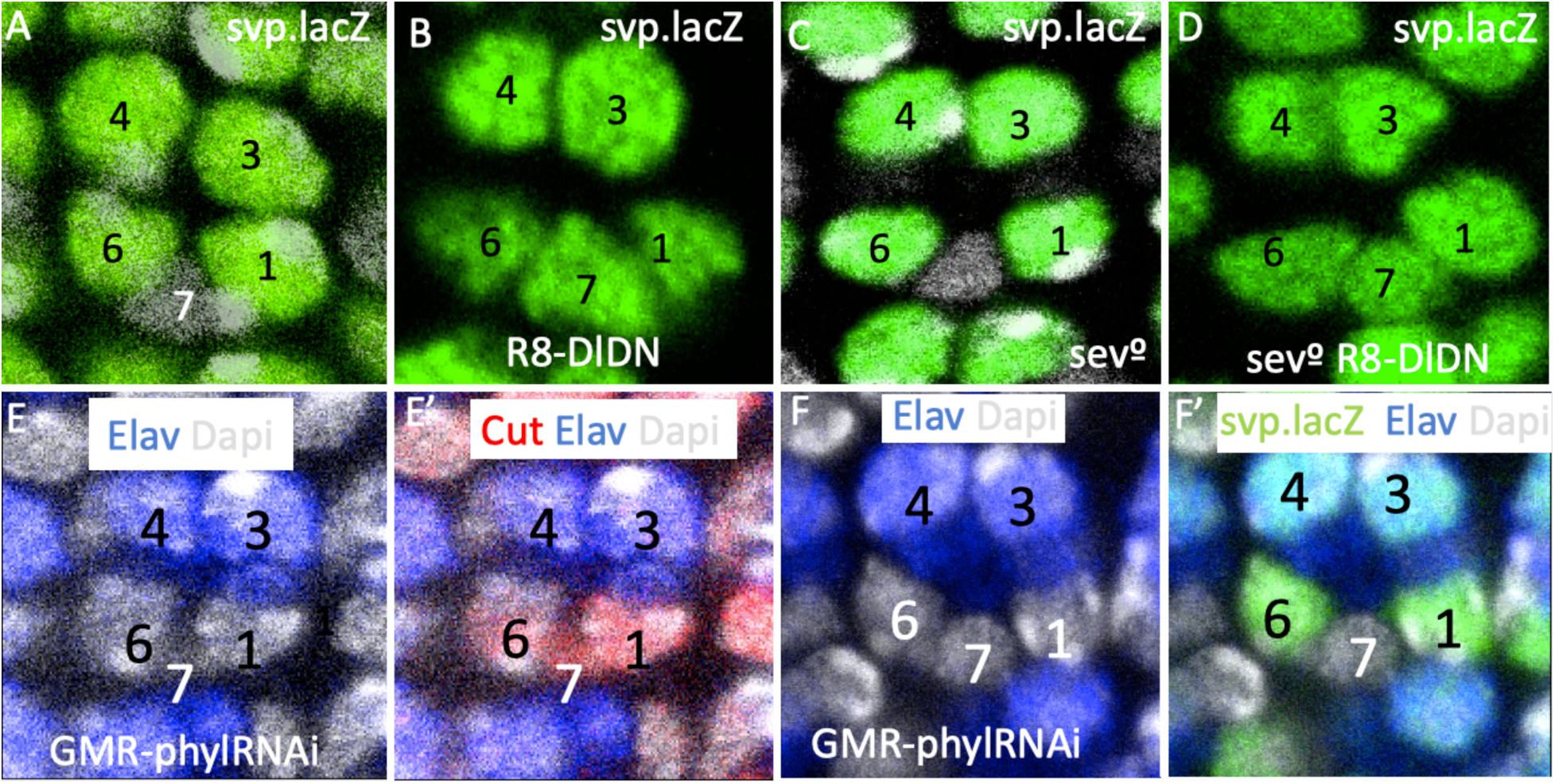
Expression of *sev.lacZ* under various genetic conditions. (**A**) In wild type eye discs, *svp.lacZ* is expressed in R3/4 and R1/6 but not R7 (labeled with Dapi). (**B**) In R8-*DlDN* discs, *svp.lacZ* is additionally expressed in the R7 precursor. (**D**) In the *sev°* mutant, there is not *svp.lacZ* expression in the R7 precursor. (**E**) In the *sev°; R8-DlDN* background, *svp.lacZ* is expressed in the R7 precursor. (**E,F**) The effects of compromising *phyl* gene function in R1/6/7 precursors. (**E/E’**) The R3/4 cells express Elav indicating their differentiation as photoreceptors, but the R/1/6/7 (labeled with Dapi) do not. Rather, they express Cut, indicating that they have adopted the cone cell fate. (**F/F’**) Only R1/6 express *svp.lacZ* when *phyl* gene function is abrogated in R1/6/7, suggesting that although all three become cone cells, a Dl signal is still sent from the R1/6 precursors to repress *sev.lacZ* expression.

In R1/6 precursors, there is no N activity, which is consistent with their expression of *svp.lacZ*. But, the question then arose as to whether the *svp* transcription in R1/6 precursors requires not only the absence of N activity but also a promotional role by the RTK pathway. One possibility here is that, in addition to N, Ttk also represses *svp* transcription. In this scenario, the RTK pathway in the R1/6 precursors clears Ttk not only to allow the cell to become a photoreceptor but also to allow the expression of Svp to ensure that the cells are not specified as R7s. To evaluate this, we examined R1/6 precursors in which *phyl* gene function is abrogated (*GMR.Gal4;UAS.phylRNAi*) and observed that these cells transformed into cone cells (as evidenced by their expression of the Cut transcription factor) and yet persisted in their expression of *svp.lacZ* (Fig.6E,F). The transformation into cone cells suggests the persistence of Ttk, and their svp.*lacZ* expression argues against a role for Ttk in preventing *svp* transcription. Interestingly, although R1/6/7 all transform into cone cells in the *GMR.Gal4 UAS.phylRNAi* background, (evidenced by their Cut expression), *svp.lacZ* expression remains repressed in the R7 precursor (Fig.6E,F). This suggests that even having undergone their fate transformation to cone cell types, the cells in the R1/6 positions still express Dl and activate N in the cell in the R7 position.

Another mechanism by which the RTK pathway could promote *svp* transcription is through its regulation of the Pnt and Yan transcription factors in a manner similar to the way *phyl* expression is controlled. Unfortunately, we lacked the tools to perform meaningful experiments in this regard and the question remains unresolved.

## Discussion

The specification of the fly R7 photoreceptor has long been a model system for understanding how fates are assigned by the different signals that cells receive, and the more we probe the molecular events that underlie the process, the more we appreciate its molecular logic. In this paper, we describe the details of two sequential decisions made by the R7 precursor. The first is to become a photoreceptor engendered by the removal of Ttk, and the second is to become an R7 rather than an R1/6 by preventing the expression of Svp. Previously, we defined three rather nebulous functions for N in R7 specification: the *anti-neural function*, the *neural-promoting function,* and the *R7-v-R1/6 function*. Now, we have a molecular understanding of these functions and how they are integrated into the decision to make an R7 photoreceptor

### Control of *phyl* transcription

Ttk acts as simple switch for the photoreceptor fate: turn it off (degrade the protein), and the photoreceptor fate is specified; leave it on (Ttk persistence), and the non-photoreceptor fate results. The switch metaphor refers to Ttk, but there is not a simple on/off state for *phyl* transcription. Rather, it is active in all R7 equivalence group cells before they are recruited to the clusters and significantly strengthens in the presumptive photoreceptors and declines in the cone cells precursors. Thus, the switch that determines the Ttk on/off state results from whether *phyl* transcriptional levels are reduced or boosted. In the R1/6 precursors, Spitz-mediated DER activation raises *phyl* transcription, but in the R7 precursor, its inherent N activity and associated *anti-neural* and *neural-promoting functions* orchestrate a more complex regulation of *phyl* expression. Previously, we identified *sev* and *dpn* as N target genes that implemented the N *neural-promoting function*, with Sev acting to hyperactivate the RTK pathway to ensure Ttk degradation and Dpn then repressing any further *ttk* transcription to prevent protein resupply. But the N target genes that mediate its *anti-neural function* were unknown. Two genes appeared as likely targets for this: *yan* and *ttk*. To evaluate the roles either might play in the N *anti-neural function*, we generated transcriptional reporters for each and observed that *yan* and not *ttk* appeared N-responsive in the R7 precursor. Specifically, we observed a burst of *yan* transcriptional activity that was lost when N signaling was abrogated in the cell. Here, we posit that the increased *yan* transcription generates extra Yan protein at a level sufficient to titrate the actions of DER-regulated MAPK to inactivate it and effectively leave a transcriptional repressor complex at the *phyl* locus.

To evaluate this model, we generated a *phyl* reporter to determine whether there was a correlating increase in *phyl* transcription when N activity was reduced in the R7 precursor. This is not what we detected; *phyl* transcriptional levels in R7 precursors with compromised N activity (*R8-DlDN*) was roughly the same as in their wild type counterparts, ostensibly suggesting that N had no effect on the gene’s expression. But in this genotype, the R7 precursor is transformed into an R1/6-like cell in which there is neither the *anti-neural* nor *neural-promoting functions*, and R1/6 levels of *phyl* transcription are expected (the same as in R7 precursors). We therefore examined *phyl* transcription *in sev°* R7 precursors, in which *the neural-promoting function* is inactive due to the lack of Sev, while the *anti-neural function* should still be operational. Here, we observed a strong decline in the *phyl* transcription. Thus, when we compare the *phyl* transcription in the *sev°* mutant background with that of *R8-DlDN*, we compare the loss of the *neural-promoting function* with the loss of both the *neural-promoting* and the *anti-neural functions* respectively. The stronger expression of the *phyl* reporter in the *R8-DlDN* background thus represents the loss of the *anti-neural function*. If this reasoning is correct, then removal of Sev from an R7 with compromised N activity (*sev°; R8-DlDN*) should have no effect on the level of *phyl* transcription because the *neural-promoting function* is already absent in R8-DLDN alone. This is what we found. We next examined what happens when we restore Sev to the R8-DLDN R7 precursor to allow the hyperactivation of the RTK pathway in the absence of the *anti-neural function*; would this drive enhanced *phyl* transcription? It did not. There are a number of possible explanations for why this did not happen, the simplest of which is that *phyl* transcription is already maximal in wild type R7s and removing the repression cannot boost it further.

#### A model for the integration of the N *anti-neural* and *neural-promoting functions*

The model we propose then is that N activity triggers an increase in *yan* transcription and that the resulting level of Yan protein is too great for DER-activated MAPK to phosphorylate and inactivate. This represents the N *anti-neural function*. N also provides the *neural-promoting* function by activating *sev* transcription. This leads to high levels of Sev protein that, when activated, trigger a hyperactivation of the RTK pathway, resulting in elevated levels of activated MAPK sufficient to negate the effects of higher Yan levels. MAPK also activates PntP2, which, in the absence of functional Yan, promotes *phyl* transcription

Developmental decisions of the R7 type frequently use redundant mechanisms and consolidation processes to ensure robust and unambiguous cell fate assignments. A useful example here is the role played by Dpn in the *neural promoting function* (Mavromatakis and Tomlinson, 2016). *dpn* transcription is promoted by N activity but is repressed by Ttk, and as RTK activity degrades that Ttk in the R7 precursor, the transcriptional repression is lost and N activity promotes *dpn* transcription. The resulting Dpn protein shuts down further transcription from the *ttk* locus, thereby preventing any resupply of Ttk protein and reinforcing the decision to become a photoreceptor. Similarly, with the *anti-neural function*, we do not expect that the increase in *yan* transcription that N activity engenders in the R7 precursor will be the sole mechanism used to oppose the specification of the cell as a photoreceptor. Indeed, our preliminary results suggest that Enhancer of Split (E(spl)) transcriptional repressor proteins, that mediate many N repressive actions, may also influence the *anti-neural function*, and we are currently working to understand if and how this happens.

### Control of *svp* transcription

Svp has long been assumed to play a key role in mediating the choice between the R7 and R1/6 fates. First, loss of Svp from R1/6 precursors transforms them into R7-like cells, suggesting that it acts to suppress the R7 fate. Second, ectopic expression of Svp in R7 precursors transforms them into R1/6-like cells, again suggesting a role in suppressing the R7 fate. Here, we demonstrate that *svp* transcription is lost in the R7 precursor when N activity is abrogated and identify it as a N target gene in the N *R7-v-R1/6 function*. This raises the question of how N activity achieves *svp* transcriptional repression. Again, we suspect the action of E(spl) proteins here, and we are currently working to determine whether they mediate the N signal that represses *svp* transcription.

Above, we have used the term *R7-like cells* to refer to the fate of R1/6 precursors in the absence of the *svp* gene. More accurately, these cells are either R7 or R8 types – both have small central rhabdomeres but different opsin expressions (Miller et al., 2008). Thus, when svp is removed from R1/6 precursors, they adopt the R7 or R8 fate, but its absence from R7 precursor invariably leads to the R7 fate, which lead Miller et al to propose yet another function for N in the R7 precursor- the role of ensuring the adoption of the R7 rather than R8 fate in the absence of Svp.

## Methods

### Generation of transcriptional reporters

For each gene, a homology vector was constructed containing four sequential DNA sequences. The first (5’) was genomic DNA from the gene beginning immediately 5 to the ATG and extending 1.5kb 5’. The second contained the coding sequence for GFP fused to MYC with a nuclear localization sequence (GFP.MYC^NLS^) followed by sv40 transcriptional termination sequences. The third contained a modified form of the *white (w)* gene coding sequence expressed under *GMR* transcriptional control (*GMR.w^+^*), flanked 3’ and 5’ by FRTs or by LoxP sequences. Fourth was a genomic fragment beginning 3’ to the most 3’ gRNA used and extending 1.5kb 3’ thereof. Three gRNAs were derived from the sequences to be deleted in the insertion were expressed in a modified pCFD3 vector (Addgene). The homology and *gRNA* vectors were co-injected into *w^-^* embryos, and the resulting adults were crossed to *w^-^* flies. Transformants were detected by the red eyes engendered by the presence of *GMR.w^+^*. The lines were balanced, and the *GMR.w^+^*cassettes were excised using Flipase or CRE expression. Each line was checked by genomic PCR to validate its site of insertion.

### Fly lines

*y^1^w*;P{GawB}sca^109-68^/CyO* FBst0006479 BDSC 6479.

*w*;P{UAS-Delta.DN}TJ2/CyO* FBstoo26697 BDSC 26697

*sev-lacZ* (Bowtell et al., 1989)

*sevd2* FBal0015458

*GMR-Gal4* (Freeman. 1996)

phylRNAi - y[1] v[1]; P{y[+t7.7] v[+t1.8]=TRiP.JF)3369 attP2 FBti0129061. BDSC 29433 svp.lacZ FBst0007314 BDSC 7314

### Immunohistochemistry and histology

Protocols for adult eye sectioning(Tomlinson and Ready, 1987) and antibody staining (Tomlinson et al., 2011) have been described previously. All antibodies used were as previously described (Mavromatakis and Tomlinson, 2016).

## Acknowledgments

We thank Jason Rudas for expert technical assistance with the Crispr transformations, Gary Struhl for comments on the manuscript, and Arianna Pepin for editing the manuscript.

